# Origin and diversification of fibroblasts from the sclerotome in zebrafish

**DOI:** 10.1101/2021.04.19.440421

**Authors:** Roger C. Ma, Katrinka M. Kocha, Emilio E. Méndez-Olivos, Tyler D. Ruel, Peng Huang

## Abstract

Fibroblasts play an important role in maintaining tissue integrity by secreting components of the extracellular matrix and initiating response to injury. Although the function of fibroblasts has been extensively studied in adults, the embryonic origin and diversification of different fibroblast subtypes during development remain largely unexplored. Using zebrafish as a model, we show that the sclerotome, a sub-compartment of the somite, is the embryonic source of multiple fibroblast subtypes including tenocytes (tendon fibroblasts), blood vessel associated fibroblasts, fin mesenchymal cells, and interstitial fibroblasts. High-resolution imaging shows that different fibroblast subtypes occupy unique anatomical locations with distinct morphologies. Long-term Cre-mediated lineage tracing reveals that the sclerotome also contributes to cells closely associated with the axial skeleton. Ablation of sclerotome progenitors results in extensive skeletal defects. Using photoconversion-based cell lineage analysis, we find that sclerotome progenitors at different dorsal-ventral and anterior-posterior positions display distinct differentiation potentials. Single-cell clonal analysis combined with in vivo imaging suggests that the sclerotome mostly contains unipotent and bipotent progenitors prior to cell migration, and the fate of their daughter cells is biased by their migration paths and relative positions. Together, our work demonstrates that the sclerotome is the embryonic source of trunk fibroblasts as well as the axial skeleton, and local signals likely contribute to the diversification of distinct fibroblast subtypes.

## INTRODUCTION

Fibroblasts are connective tissue cells present in most organs of animals. They function as tissue support cells by synthesizing and remodeling the extracellular matrix (ECM). Recent studies have shown that tissue-resident fibroblasts also play important regulatory roles in wound healing, inflammation, tumor microenvironment, and tissue fibrosis (Kalluri, 2016; Kendall and Feghali-Bostwick, 2014; Plikus et al., 2021). Although fibroblasts have historically been viewed as a homogenous cell population, recent work using single-cell RNA sequencing (scRNA-seq) has revealed a high level of heterogeneity in fibroblast populations from different tissues as well as within the same tissue (Buechler et al., 2021; Forte et al., 2022; Muhl et al., 2020; Tabib et al., 2018; Xie et al., 2018). This raises the question of how the different fibroblast subtypes are generated during embryonic development.

One mechanism to diversify fibroblast populations is through contributions from multiple embryonic sources. Previous studies suggest that fibroblasts in the same tissue may have heterogeneous embryonic origins. For example, in avian and mouse embryos, dermal fibroblasts in the head are of neural crest origin, whereas those in the dorsal and ventral trunk originate from the somite and lateral plate mesoderm, respectively (Thulabandu et al., 2018). Similarly, during heart development in chick and mouse embryos, most cardiac fibroblasts originate from lateral plate mesoderm derived epicardium, while a small population arises from an endothelial/endocardial lineage (Tallquist, 2020).

Somites are the embryonic source of many tissue support cells, including fibroblasts (Christ et al., 2007). During development, somite forms three separate compartments: the dermatome, myotome, and sclerotome. The sclerotome contributes to the axial skeleton of animals (Christ et al., 2004). Work in mouse and chick has shown that the sclerotome is also the developmental origin of tendon fibroblasts (tenocytes) (Brent et al., 2003; Schweitzer et al., 2001) and vascular smooth muscle cells (Pouget et al., 2008; Wiegreffe et al., 2007). In zebrafish, the sclerotome has a unique bipartite organization and contributes to tenocytes, perivascular fibroblasts, and pericytes in a stereotypical manner (Ma et al., 2018; Rajan et al., 2020). It raises the question of the cellular mechanisms governing the diversification of fibroblast subtypes from sclerotome progenitors.

In fish and amphibians, fin mesenchymal cells are a population of fibroblasts present in the larval fin fold. They express many ECM proteins to provide structural support for the developing fin fold (Durán et al., 2011; Feitosa et al., 2012). However, the developmental origin of fin mesenchymal cells remains controversial. Dye labeling experiments in *Xenopus* and zebrafish suggest that the neural crest contributes to the fin mesenchyme (Smith and Hall, 1990; Smith et al., 1994). However, later experiments in *Xenopus* suggest that the mesoderm also contributes to fin mesenchymal cells in both the dorsal and ventral fins (Garriock and Krieg, 2007; Tucker and Slack, 2004). Similarly, cell transplantation experiments in axolotls reveal dual origin of the fin mesenchyme from both the neural crest and somites (Sobkow et al., 2006). More recent work using genetic lineage tracing in fish (zebrafish) and amphibians (*Xenopus* and axolotl) supports a model where fin mesenchymal cells originate exclusively from the mesoderm with no contribution from the neural crest lineage (Lee et al., 2013; Taniguchi et al., 2015). In particular, the dermomyotome compartment of the somite has been suggested as the source of fin mesenchymal cells in zebrafish (Lee et al., 2013).

Here, we show that the sclerotome in zebrafish is a major source of multiple fibroblast populations as well as cells associated with the axial skeleton. Different sclerotome-derived fibroblast subtypes display unique morphologies and occupy distinct anatomical locations. In vivo time-lapse imaging and single-cell clonal analysis reveal that sclerotome progenitors can generate multiple fibroblast subtypes, and their differentiation potential is biased by their positions as well as their migratory patterns. Together, our results suggest that distinct fibroblast subtypes are specified from sclerotome progenitors by local cellular environments.

## RESULTS

### Characterization of a sclerotome reporter in zebrafish

To explore the lineage potential of the sclerotome in zebrafish, we recently developed a sclerotome reporter, *nkx3.1:Gal4; UAS:NTR-mCherry* (*nkx3.1*^*NTR-mCherry*^) (Ma et al., 2018). The *nkx3.1:Gal4* driver accurately recapitulated the expression of the endogenous *nkx3.1* gene (Fig S1). The specificity of this *nkx3.1*^*NTR-mCherry*^ line was further validated by co-labeling with known sclerotome markers, including *vcanb* (*versican b*) (Landolt et al., 1995), *foxc1b* (Topczewska et al., 2001), and *twist1b* (previously known as *twist1a*) (Germanguz et al., 2007; Yeo et al., 2009) (Fig 1A). Due to the perdurance of the mCherry protein, we used *nkx3.1*^*NTR-mCherry*^ to label the initial sclerotome domains and all their descendants. We crossed *nkx3.1*^*NTR-mCherry*^ with the endothelial reporter *kdrl:EGFP* to examine sclerotome-derived cells at 2 dpf (days post-fertilization). Based on the anatomical location and morphology, we broadly defined five groups of mCherry^+^ cells from the *nkx3.1* lineage (Fig 1B-C): 1) cells closely associated with blood vessels in the trunk, including the dorsal longitudinal anastomotic vessel (DLAV), intersegmental vessels (ISV), dorsal aorta (DA), and caudal vein plexus (CVP); 2) tenocytes along somite boundaries as previously described (Ma et al., 2018; Subramanian et al., 2018); 3) interstitial cells located in the space between the notochord, spinal cord, and muscles; 4) cells located in both dorsal and ventral fin folds; and 5) some muscle cells located at the dorsal and ventral edges of each somite (Ma et al., 2018). The contribution of muscles from the *nkx3.1* lineage suggests that *nkx3.1* labels some early bipotent somitic cells that generate both sclerotome progenitors and muscles, which is consistent with previous single-cell lineage analysis of the ventral somite (Morin-Kensicki and Eisen, 1997). In addition to the five groups of cells from the *nkx3.1* lineage, the *nkx3.1:Gal4* line drove ectopic expression in some cuboidal cells at the edge of the fin fold (Fig 1C and S2), which were not derived from the sclerotome. It is worth noting that although the *nkx3.1:Gal4* line nicely matched the endogenous *nkx3.1* expression, the amplification of the GAL4/UAS system in *nkx3.1*^*Kaede*^ (*nkx3.1:Gal4; UAS:Kaede*) and *nkx3.1*^*NTR-mCherry*^ lines resulted in various degrees of ectopic expression outside the *nkx3.1* domains (Fig S1). We have previously shown that both tenocytes and ISV-associated perivascular fibroblasts express *col1a2* and *col5a1* (Ma et al., 2018; Rajan et al., 2020), two pan-fibroblast markers defined by scRNA-seq experiments (Metikala et al., 2021; Muhl et al., 2020). Indeed, all *nkx3.1*^*NTR-mCherry*^-positive non-muscle cells (groups 1-4) were labeled by *col1a2:GFP* at 2 dpf (Fig 1D), confirming their fibroblast identity. Our results suggest that the sclerotome is the embryonic source of multiple types of fibroblasts in zebrafish.

**Fig 1.**
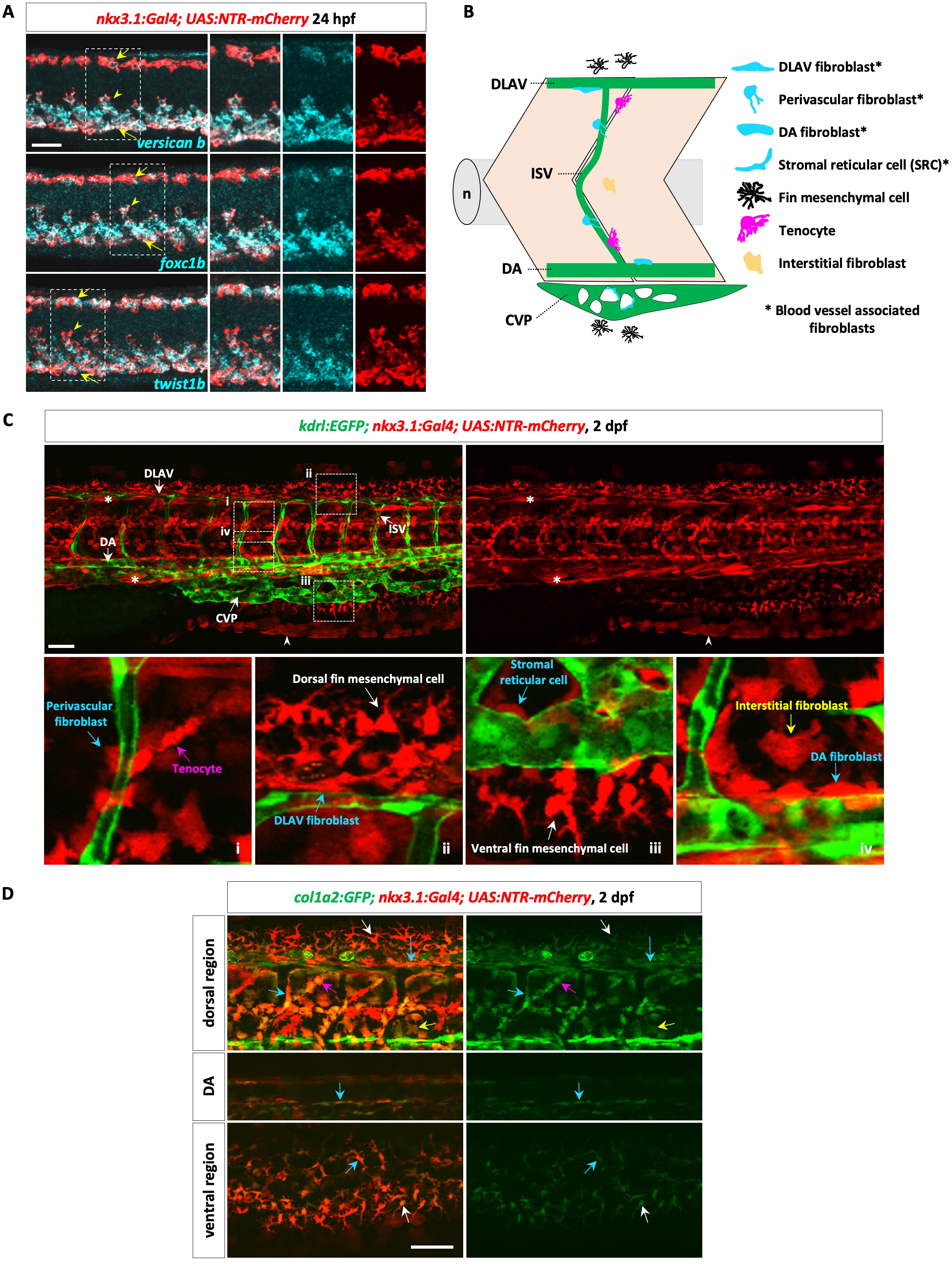
The sclerotome gives rise to different types of fibroblasts. (A) Double fluorescent in situ staining of *nkx3.1*^*NTR-mCherry*^ embryos at 24 hpf showed co-expression of the sclerotome markers *versican b, foxc1b*, and *twist1b* (cyan) with *ntr-mcherry* (red) in the dorsal sclerotome (short arrows), ventral sclerotome (long arrows), and sclerotome-derived notochord-associated cells (arrowheads). *n*=15 per staining. (B) Schematic of sclerotome-derived fibroblasts: blood vessel associated fibroblasts (cyan), fin mesenchymal cells (black), tenocytes (magenta), and interstitial fibroblasts (yellow). DLAV, dorsal longitudinal anastomotic vessel; ISV, intersegmental vessel; DA, dorsal aorta; CVP, caudal vein plexus; n, notochord. (C) At 2 dpf, *nkx3.1*^*NTR-mCherry*^-positive cells (red) were observed throughout the trunk showing close association with the vasculature labeled by *kdrl:EGFP* (green) (top panels). Zoomed-in views (bottom panels) show different mCherry^+^ fibroblast subtypes (arrows) in four boxed regions of the trunk (i-iv). mCherry expression in muscles (asterisks) and skin cells along the fin fold (arrowheads) is shown. *n*=13. (D) Most *nkx3.1*^*NTR-mCherry*^-positive sclerotome-derived cells (red) co-expressed *col1a2:GFP* (green) at 2 dpf. The upper panel shows the dorsal region with dorsal fin mesenchymal cells (white arrows), DLAV fibroblasts (long cyan arrows), ISV-associated perivascular fibroblasts (cyan arrows), tenocytes (magenta arrows), and interstitial fibroblasts (yellow arrows). The middle panel shows DA fibroblasts (cyan arrows) associated with the dorsal aorta. The bottom panel shows stromal reticular cells (cyan arrows) and ventral fin mesenchymal cells (white arrows). *n*=12. Scale bars: 50 μm.

### The sclerotome generates different types of trunk fibroblasts

We performed mosaic labeling and marker analysis to characterize fibroblasts derived from the *nkx3.1* lineage (Fig 2A-C). *nkx3.1*^*NTR-mCherry*^-expressing fibroblasts in the trunk, including tenocytes, blood vessel associated fibroblasts, and interstitial fibroblasts, were marked by pan-fibroblast markers such as *tgfbi* and *col5a1* (Metikala et al., 2021; Muhl et al., 2020) (Fig 2B). A subset of these sclerotome-derived fibroblasts was associated with different vascular beds (Fig 1C and 2A). We previously described ISV-associated perivascular fibroblasts, characterized by a globular cell body with short processes extending around the ISV (Rajan et al., 2020). By contrast, labeling with a mosaic *col1a2:Gal4; UAS:Kaede* line (*col1a2*^*Kaede*^) (Sharma et al., 2019) revealed that fibroblasts associated with DLAV or DA (DLAV and DA fibroblasts, respectively) were more elongated along the long axis of the blood vessel (Fig 1C and 2A). CVP is a transient venous network with a stereotypical “honeycomb-like” structure. Fibroblasts marked by *nkx3.1*^*NTR-mCherry*^ occupied the space in the honeycomb pockets of the CVP (Fig 1C and 2A). Mosaic labeling showed that individual Kaede^+^ cells had a small and elongated cell body with long cellular processes along the underlying endothelium and sometimes across the honeycomb pocket (Fig 2A). The location and morphology of these CVP-associated fibroblasts suggest that they are stromal reticular cells (SRCs) (Murayama et al., 2015).

**Fig 2.**
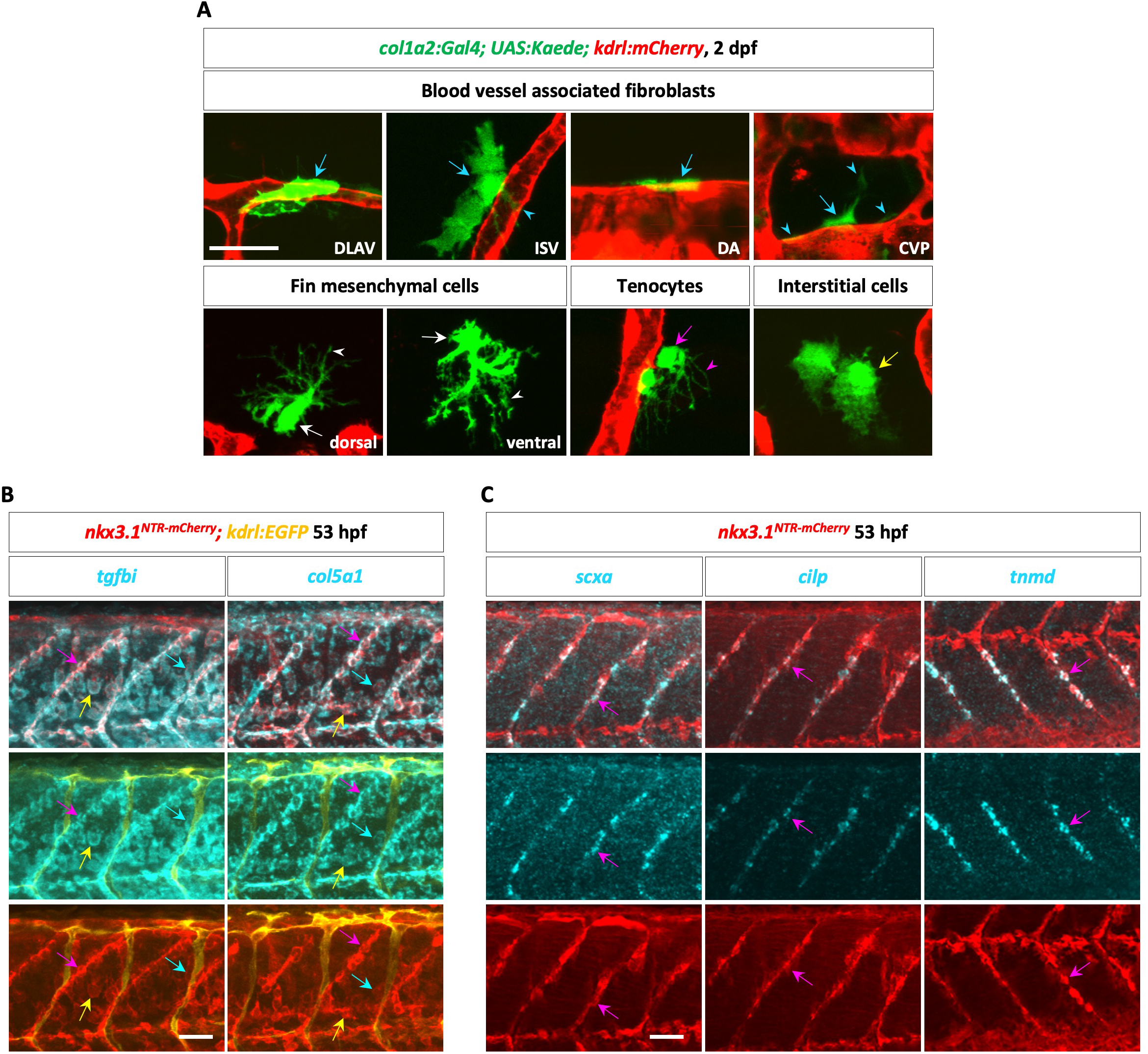
Characterization of sclerotome-derived fibroblasts. (A) The mosaic *col1a2*^*Kaede*^ line (green) was crossed with *kdrl:mCherry* (red) to visualize the morphology of different fibroblasts at a high resolution at 2 dpf: blood vessel associated fibroblasts (cyan arrows), fin mesenchymal cells (white arrows), tenocytes (magenta arrow), and interstitial fibroblasts (yellow arrow). Cell processes are indicated by arrowheads. The associated vascular bed and fin fold are indicated for blood vessel associated fibroblasts and fin mesenchymal cells, respectively. *n*=20. (B) *nkx3.1*^*NTR-mCherry*^; *kdrl:EGFP* embryos at 53 hpf were co-stained with RFP antibody (red), GFP antibody (yellow), and probes for fibroblast markers *tgfbi* or *col5a1* (cyan). mCherry^+^ blood vessel associated fibroblasts (cyan arrows), tenocytes (magenta arrows), and interstitial fibroblasts (yellow arrows) co-expressed *tgfbi* and *col5a1. n*=12 per staining. (C) *nkx3.1*^*NTR-mCherry*^ embryos at 53 hpf were co-stained with RFP antibody (red) and probes for tenocyte markers *scxa, cilp*, or *tnmd* (cyan). mCherry^+^ tenocytes (magenta arrows) along MTJ co-expressed tenocyte markers. *n*=12 per staining. Scale bars: (A) 20 μm; (B, C) 50 μm.

Tenocytes and interstitial fibroblasts were the two fibroblast subtypes that populated the trunk region but were not closely associated with blood vessels (Fig 1C). We and others have previously shown that tenocytes are located along the myotendinous junction (MTJ), extending long cellular processes into the intersomitic space (Fig 2A) (Ma et al., 2018; Subramanian et al., 2018). Marker analysis revealed that *nkx3.1*^*NTR-mCherry*^-positive cells along the MTJ were specifically labeled by known tendon markers such as *scxa, cilp*, and *tnmd* (Chen and Galloway, 2014; Ralliere et al., 2015) (Fig 2C). By contrast, interstitial fibroblasts were scattered throughout the interstitial space and displayed varied morphologies (Fig 1C and 2A).

### The sclerotome contributes to fin mesenchymal cells

The *nkx3.1*^*NTR-mCherry*^ line labeled cells in the fin fold (Fig 1B-C). At 2 dpf, the fin fold is subdivided into two parts: the major lobe wrapping around the trunk and tail, and the minor lobe beneath the yolk extension (Parichy et al., 2009) (Fig S2). Cells from the *nkx3.1* lineage populated both the dorsal and ventral regions in the major lobe (Fig 1C and S2). Using the mosaic *col1a2*^*Kaede*^ line, we found that Kaede^+^ fin cells displayed extensive “tree-like” cellular processes projecting towards the periphery of the fin fold (Fig 2A). The location and morphology suggest that these sclerotome-derived fin fold residing cells correspond to fin mesenchymal cells (Carney et al., 2010; Feitosa et al., 2012; Lee et al., 2013). Previous studies have shown that fin mesenchymal cells are marked by *hemicentin2* (*hmcn2*) and *fibulin1* (*fbln1*), while the neighboring cell layer of apical epidermal cells is labelled by *hmcn2* and *fras1* (Carney et al., 2010; Feitosa et al., 2012; Lee et al., 2013) (Fig 3A). Double staining revealed two juxtaposing cell layers in the fin fold: *hmcn2*^*+*^*fbln1*^*+*^ fin mesenchymal cells in the proximal layer, and *hmcn2*^*+*^*fras1*^*+*^ apical epidermal cells in the distal layer (Fig 3B). As expected, double staining in *nkx3.1*^*NTR-mCherry*^ embryos showed that mCherry^+^ cells in the fin fold co-expressed *fbln1* but not *fras1* (Fig 3C), confirming that the sclerotome contributes to fin mesenchymal cells, a specific cell population in the fin fold.

**Fig 3.**
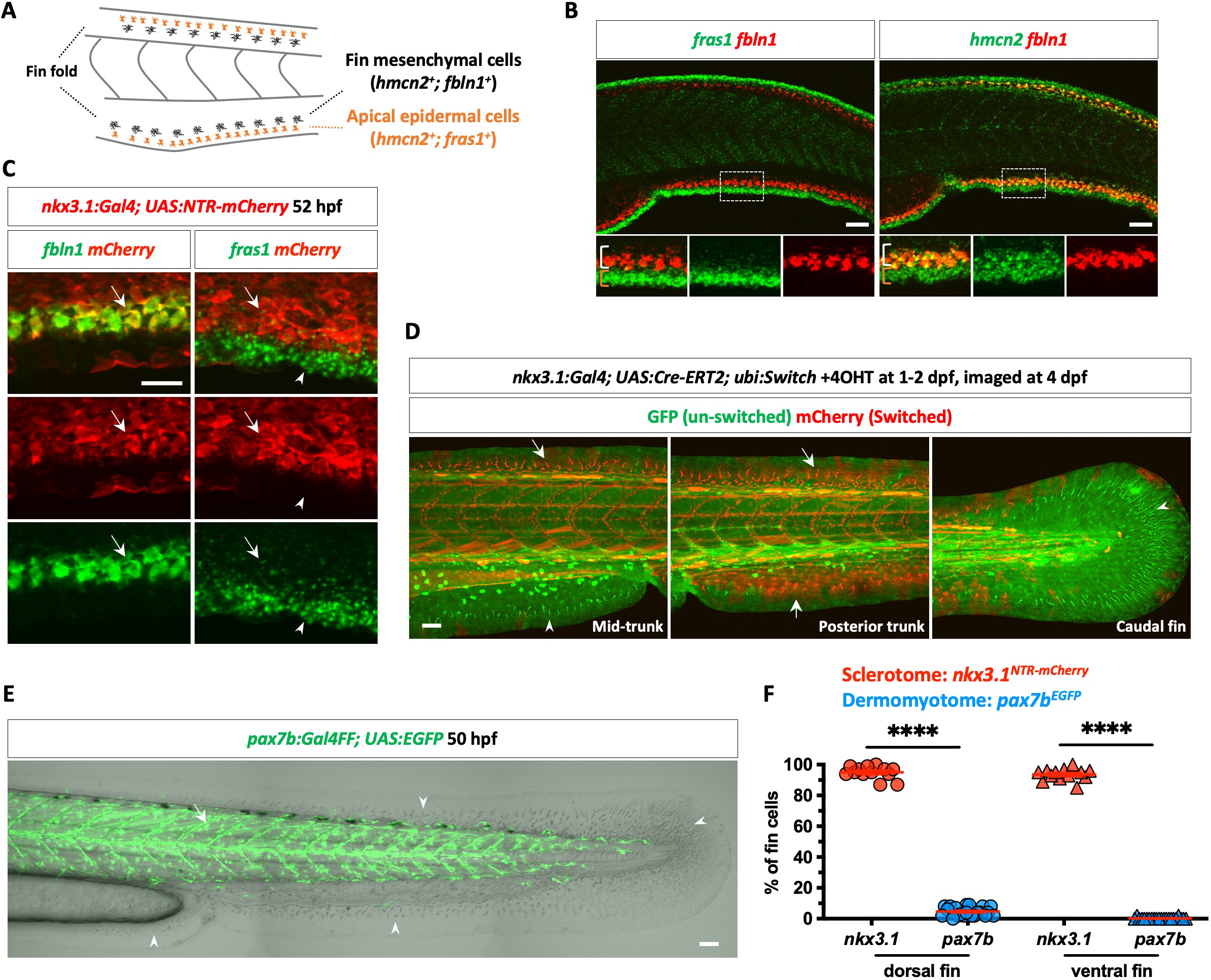
The sclerotome contributes to fin mesenchymal cells. (A) Schematic of the zebrafish fin fold. Both dorsal and ventral fin folds are populated by two adjacent cell layers: the distal *hmcn2*^*+*^*fras1*^*+*^ apical epidermal cells and proximal *hmcn2*^*+*^*fbln1*^*+*^ fin mesenchymal cells. (B) Double fluorescent in situ staining of *fbln1* (red) with either *fras1* or *hmcn2* (green) at 48 hpf. *fras1* labeled apical epidermal cells (orange brackets), *fbln1* marked fin mesenchymal cells (white brackets), while *hmcn2* was expressed in both layers. *n*=15 per staining. (C) *nkx3.1*^*NTR-mCherry*^ embryos were costained with *fbln1* or *fras1* probes (green) and anti-RFP antibody (red) at 52 hpf. Apical epidermal cells and fin mesenchymal cells are indicated by arrowheads and arrows, respectively. *n*=10 per staining. (D) *nkx3.1*^*Cre-ERT2*^; *ubi:Switch* embryos treated with 4OHT at 1-2 dpf and imaged at 4 dpf. Switched mCherry^+^ fin mesenchymal cells were present in the dorsal and ventral regions of the major lobe (arrows) but absent in the minor lobe or caudal fin (arrowheads). *n*=10. (E) *pax7b*^*EGFP*^ embryos at 50 hpf showed extensive EGFP expression in the dermomyotome (arrow), while most fin mesenchymal cells (arrowheads) in both the major and minor lobes of the fin fold were EGFP^-^. *n*=10. (F) Quantification of lineage contribution of fin mesenchymal cells. The percentage of fin mesenchymal cells labeled by each lineage reporter at 2 dpf was plotted as mean±SEM. *n*=22 (*pax7b*^*EGFP*^) and 13 (*nkx3.1*^*NTR-mCherry*^). Statistics: Mann-Whitney *U* test; p<0.0001 (****). Scale bars: (B, D, E) 50 μm; (C) 20 μm.

Interestingly, *nkx3.1*^*Kaede*^-expressing fin mesenchymal cells were absent in the caudal fin fold and minor lobe (Fig S2). Cre-mediated lineage tracing in *nkx3.1:Gal4; UAS:Cre-ERT2; ubi:Switch* fish showed similar results (Fig 3D). Switched mCherry^+^ cells derived from the *nkx3.1* lineage gave rise to fin mesenchymal cells in both the dorsal and ventral regions of the major lobe at 4 dpf, but cells in the minor lobe and caudal fin remained GFP^+^mCherry^-^ (un-switched) (Fig 3D). Our results suggest that sclerotome is not the only lineage that gives rise to fin mesenchymal cells. To determine whether the dermomyotome (also known as the external cell layer) contributes to fin mesenchymal cells, we imaged *pax7b:Gal4FF; UAS:EGFP* (*pax7b*^*EGFP*^) fish at 50 hpf. Interestingly, despite extensive labeling of dermomyotome cells in all somites, most fin mesenchymal cells were not marked by *pax7b*^*EGFP*^ (Fig 3E). EGFP^+^ fin cells were not observed in the minor lobe or caudal fin, while on average, only 4.6% and 0.2% of fin mesenchymal cells were *pax7b*^*EGFP*^-positive in the dorsal and ventral regions of the major lobe, respectively (Fig 3F). This is in striking contrast to 95% (dorsal) and 94% (ventral) contributions from the *nkx3.1*^*NTR-mCherry*^ reporter (Fig 3F). Therefore, our results suggest that the sclerotome, but not the dermomyotome, is the main contributor of fin mesenchymal cells in the dorsal and ventral fin folds.

Together, our data show that the sclerotome generates multiple types of fibroblasts in the zebrafish trunk, supporting different tissues such as muscles, blood vessels, and the fin fold. These fibroblast subtypes reside in distinct anatomical locations with unique morphologies, some of which can be defined by specific marker expression.

### The sclerotome contributes to the axial skeleton in zebrafish

Since the sclerotome is known to contribute to the axial skeleton (Christ et al., 2004), we performed Cre-mediated lineage tracing on cells from the *nkx3.1* lineage in zebrafish. Sclerotome-derived cells in *nkx3.1:Gal4; UAS:Cre-ERT2; ubi:Switch* fish were switched to express mCherry with 4-hydroxytamoxifen (4OHT) treatment at 1-2 dpf. Co-labeling of calcein, a vital dye for calcified structures, in switched fish at 17 dpf revealed that mCherry^+^ cells were closely associated with the vertebrae, including the centrum, neural arch, and hemal arch (Fig 4A). Interestingly, mCherry^+^ cells formed a thin layer external to the calcein-positive centrum, with enrichment in the region between adjacent centra (Fig 4A). By contrast, calcein staining in neural and hemal arches appeared to be embedded within mCherry^+^ cells with overlapping staining (Fig 4A). Long-term tracing of *nkx3.1*^*+*^ cells resulted in substantial labeling of tenocytes at the MTJ and cells associated with the vertebral column in fish at 6 weeks post-fertilization (wpf) and adult stages (Fig 4B). To determine the function of sclerotome-derived cells in skeletal development, we performed genetic ablation using the nitroreductase (NTR) system (Curado et al., 2008). *nkx3.1*^*NTR-mCherry*^ embryos or their mCherry^-^ siblings were treated with metronidazole (MTZ) at 1-2 dpf and stained for bone and cartilage at 3 wpf (Fig 4C-E). Since many ablated fish developed heart edema, likely caused by the depletion of perivascular fibroblasts (Rajan et al., 2020), we can only analyze survivors from the ablated group, which likely represents fish with more mosaic expression. MTZ-treated mCherry^+^ fish were significantly smaller than mCherry^-^ controls (Fig 4D), suggesting impaired growth with partial ablation of the *nkx3.1* lineage. Interestingly, while vertebral bodies formed normally, both neural and hemal arches displayed variable defects, with an average of 35% of vertebrae missing neural arches and 3% missing hemal arches (Fig 4C-E). Together, our results suggest that sclerotome-derived cells in zebrafish contribute to the normal development of the vertebral column, especially skeletal elements attached to the vertebral body.

**Fig 4.**
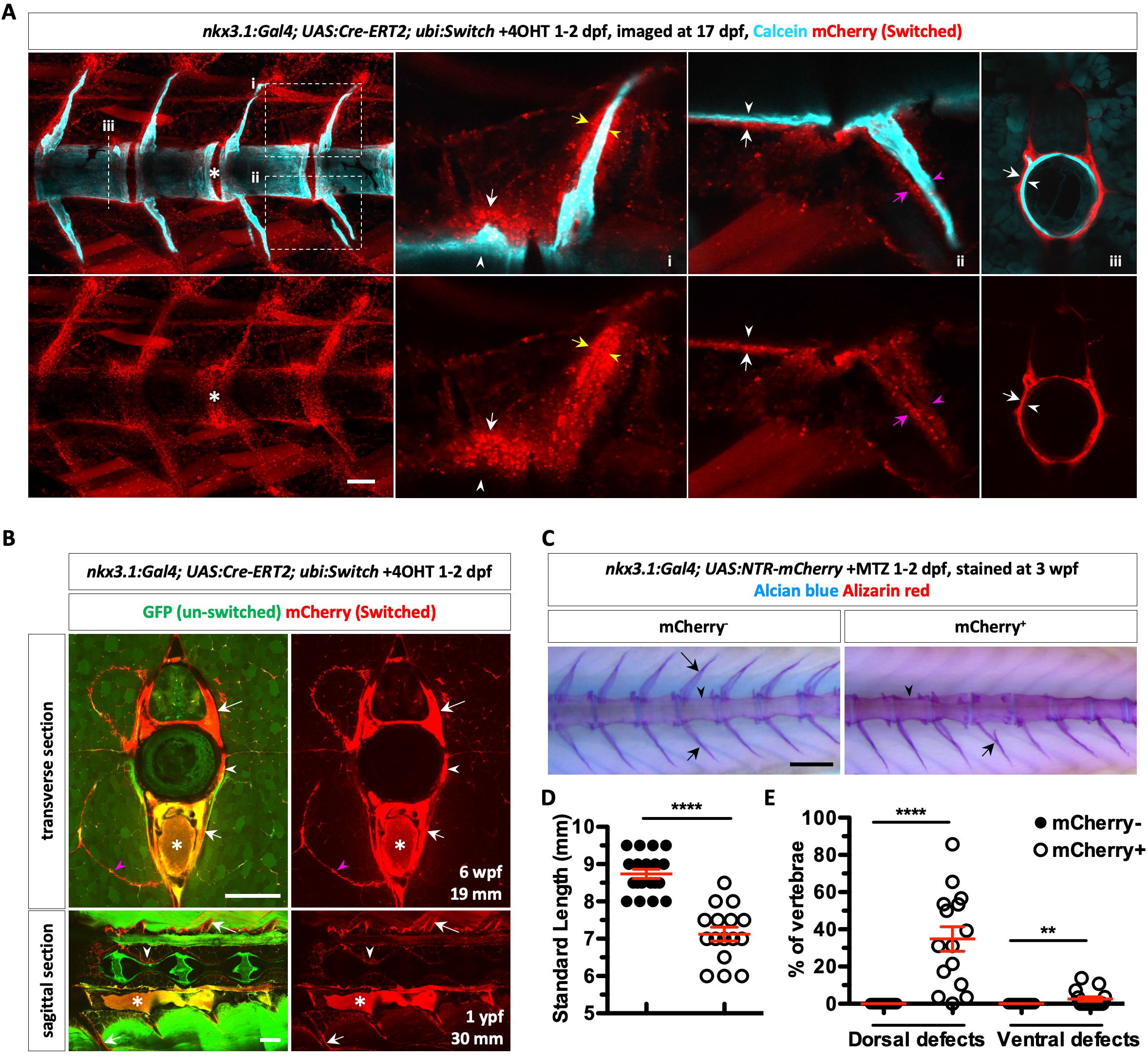
The sclerotome contributes to the axial skeleton. (A) *nkx3.1*^*Cre-ERT2*^; *ubi:Switch* embryos were treated with 4OHT at 1-2 dpf to switch on mCherry expression (red) in sclerotome-derived cells. Switched embryos were stained with calcein (cyan) at 17 dpf to visualize calcified structures, including the centra (white arrowheads), neural arches (yellow arrowheads), and hemal arches (magenta arrowheads). Magnified views of the neural (i) and hemal (ii) arches as well as a cross-sectional view of the centrum (dotted line, iii) are shown. Switched mCherry^+^ cells associated with different skeletal structures are denoted by arrows with the corresponding colors. Enrichment of mCherry^+^ cells in the regions between the vertebral bodies is indicated by asterisks. Since the calcein signal is very strong, un-switched GFP expression is undetectable under the imaging setting. *n*=5. (B) *nkx3.1*^*Cre-ERT2*^; *ubi:Switch* embryos treated with 4OHT at 1-2 dpf were raised and imaged at 6 wpf (transverse section) and 1-year post-fertilization (ypf) (sagittal section). mCherry^+^ cells were associated with the MTJ (magenta arrowheads), centrum (white arrowheads), neural arch (long arrows), and hemal arch (short arrows). The dorsal aorta (asterisks) showed strong autofluorescence in both channels. Standard lengths of imaged fish are indicated. *n*=2 per stage. (C) *nkx3.1*^*NTR-mCherry*^ embryos and their mCherry^-^ siblings were treated with MTZ at 1-2 dpf and stained with alizarin red and alcian blue at 3 wpf. mCherry^+^ fish showed normal vertebral bodies (arrowheads) with missing neural arches (long arrow) and some misconnected hemal arches (short arrows). *n*=19 (mCherry^-^) and 15 (mCherry^+^). (D, E) Quantification of standard length and skeletal defects in MTZ-treated fish (C). The percentage of vertebrae with missing neural or hemal arches was plotted as mean±SEM. Statistics: Mann-Whitney *U* test. Asterisk representation: p<0.01 (**), p<0.0001 (****). Scale bars: (A) 50 μm; (B, C) 200 μm.

### Both dorsal and ventral sclerotome domains give rise to multiple types of fibroblasts

We have previously shown that the zebrafish sclerotome consists of a dorsal and ventral domain, both of which generate tenocytes and perivascular fibroblasts (Ma et al., 2018; Rajan et al., 2020). We asked how each sclerotome domain contributes to different fibroblast subtypes. Taking advantage of the photoconvertible Kaede protein, we performed lineage tracing of sclerotome domains in *nkx3.1*^*Kaede*^ fish (Fig 5A). Either the dorsal or ventral sclerotome domain of a single somite was photoconverted from Kaede^green^ to Kaede^red^ at 24 hpf (hours post-fertilization). We then determined the fate of Kaede^red^ cells after 24 hours based on their location and morphology.

**Fig 5.**
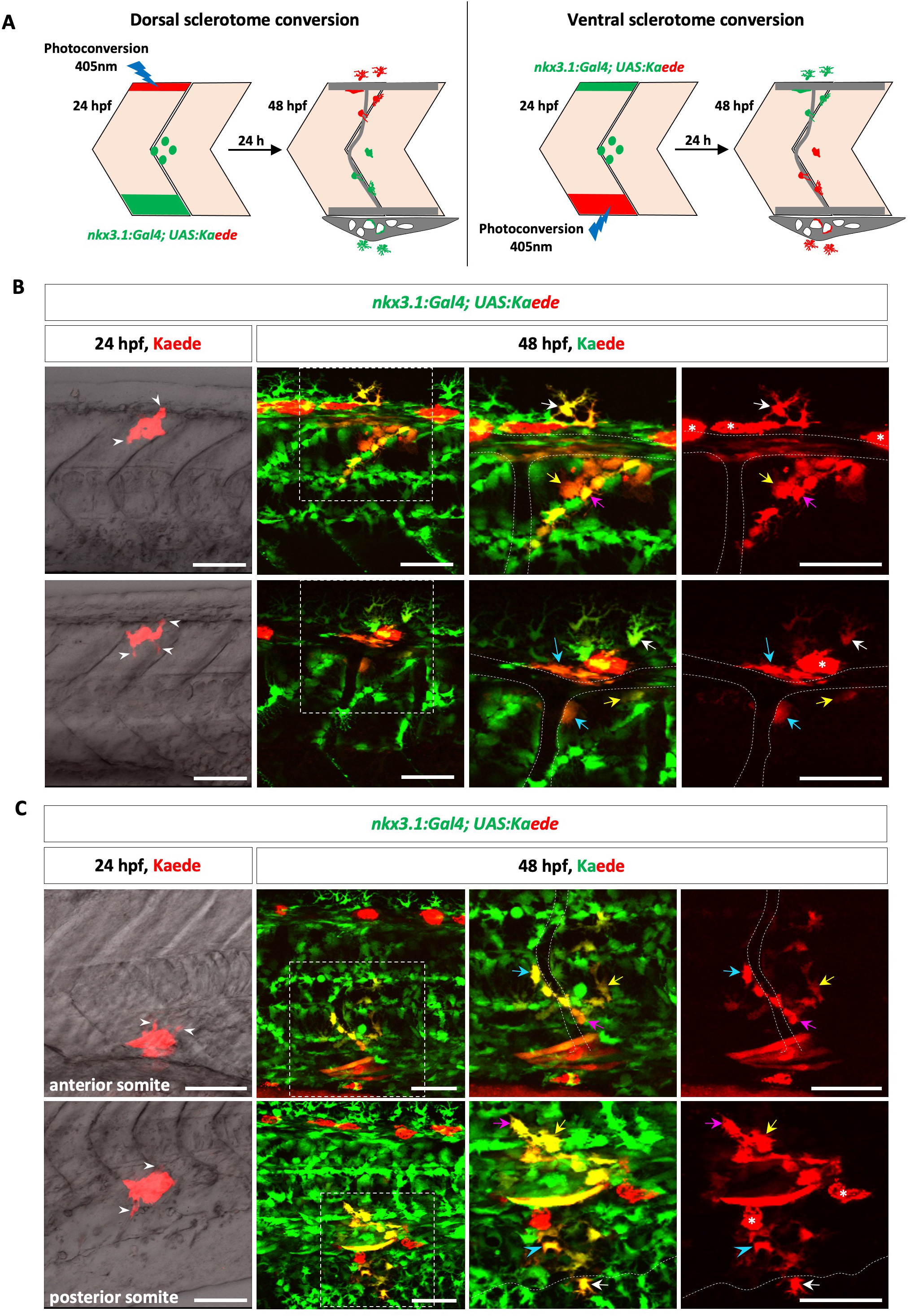
Lineage analysis of sclerotome domains. (A) Schematic of lineage analysis. The entire dorsal or ventral sclerotome domain in *nkx3.1*^*Kaede*^ embryos was photoconverted at 24 hpf and analyzed for cell composition at 48 hpf. (B, C) Examples of cell tracing of the dorsal (B) and ventral (C) sclerotome domains in *nkx3.1*^*Kaede*^ embryos. At 48 hpf, full view of the trunk and close-up views of the boxed regions are shown. Cell projections at 24 hpf (arrowheads) and the outline of blood vessels (dotted lines) are denoted. Pigment cells (asterisks) showed strong autofluorescence in the red channel. (B) Dorsal sclerotome domain generated dorsal fin mesenchymal cells (white arrows), DLAV fibroblasts (long cyan arrows), perivascular fibroblasts (cyan arrows), tenocytes (magenta arrows), and interstitial fibroblasts (yellow arrows). *n*=20. (C) In the anterior somites above the yolk extension (upper panel), the ventral sclerotome domain generated perivascular fibroblasts (cyan arrows), tenocytes (magenta arrows), and interstitial fibroblasts (yellow arrows). In the posterior somites above the CVP region (bottom panel), the ventral sclerotome domain generated SRCs (cyan arrowheads), ventral fin mesenchymal cells (white arrows), tenocytes (magenta arrows), and interstitial fibroblasts (yellow arrows). The ventral boundary of the CVP is indicated by dotted lines. *n*=20. Scale bars: 50 μm.

When the dorsal sclerotome was labeled at 24 hpf, we observed small projections from these Kaede^red^ cells, suggesting the initiation of cell migration at this stage (Fig 5B). At 48 hpf, all dorsal sclerotome-derived Kaede^red^ cells were restricted to the upper half of the trunk dorsal to the notochord (Fig 5B). Based on the morphology and location relative to the vasculature, we determined that the dorsal sclerotome generated dorsal fin mesenchymal cells, DLAV fibroblasts, tenocytes, ISV-associated perivascular fibroblasts, and interstitial fibroblasts.

Next, we performed photoconversion experiments to mark the ventral sclerotome with Kaede^red^ at 24 hpf. Similar to the dorsal domain, small cellular projections were visible from Kaede^red^ cells, suggesting that both dorsal and ventral sclerotome progenitors initiate their migration at the same stage (Fig 5C). Interestingly, in anteriorly converted somites (somites 16-18), cellular projections pointed exclusively in the dorsal direction, while in posteriorly converted somites (somites 23-25), both dorsal and ventral projections were observed (Fig 5C). This result suggests that the migration pattern is different in ventral sclerotome progenitors along the anterior-posterior axis, which likely accounts for the cells generated in the CVP and ventral fin fold region. All the cells generated by the ventral sclerotome were restricted to approximately the ventral two-thirds of the trunk. In anterior somites, the ventral sclerotome generated cells in the trunk, including tenocytes, perivascular fibroblasts, and interstitial fibroblasts at 48 hpf (Fig 5C). In contrast, the ventral sclerotome of posterior somites gave rise to cells in the trunk and cells ventral to the photoconverted region in the periphery, including CVP-associated SRCs and fin mesenchymal cells in the ventral fin fold (Fig 5C).

Together, our results suggest that both the dorsal and ventral sclerotome domains can generate multiple fibroblast subtypes, but the progenitors in each domain generate an anatomically restricted population of cells according to their dorsal-ventral and anterior-posterior positions.

### Fate of dorsal sclerotome progenitors is biased by the migration direction

Both dorsal and ventral sclerotome contribute to multiple types of fibroblasts. It is possible that sclerotome progenitors are multipotent, or alternatively, individual progenitors are “fate-restricted” to differentiate into a specific fibroblast subtype. To distinguish these possibilities, we performed single-cell clonal analysis of sclerotome progenitors as previously described (Sharma et al., 2019). We focused on the smaller dorsal domain to ensure precise single-cell photoconversion and accurate cell tracing. We selected *nkx3.1*^*Kaede*^ fish with more mosaic Kaede expression to minimize the photoconversion of multiple cells. At 24 hpf, a single Kaede^green^ cell in the dorsal domain between somites 16 and 18 was photoconverted to Kaede^red^ (Fig 6A). We then imaged the same fish after 24 hours to determine the identity of their Kaede^red^ descendants. Most clones (35/42, 83%) contained at least 2 daughter cells, suggesting at least one cell division between 24 and 48 hpf (Fig 6B). From 42 dorsal sclerotome progenitors, 38% (16/42) produced clones with at least two fibroblast subtypes, and 2% (1/42) generated three subtypes (Fig 6C). This result suggests that some dorsal sclerotome progenitors are bipotent and possibly multipotent at 24 hpf. Consistent with domain tracing, we found that dorsal sclerotome progenitors gave rise to all four types of fibroblasts, including dorsal fin mesenchymal cells, tenocytes, blood vessel associated fibroblasts, and interstitial fibroblasts (Fig 6D). As observed earlier, 33 of the 42 photoconverted Kaede^red^ cells at 24 hpf showed visibly biased projections pointing either dorsally (14 cells) or ventrally (19 cells), likely corresponding to their migration direction. To correlate progenitor morphology with their lineage potential, we categorized different fibroblasts into two types based on their final positions (Fig 6A). In the type I group, fibroblasts are located dorsal to the somite, including dorsal fin mesenchymal cells and DLAV fibroblasts. Type II cells, such as tenocytes, ISV-associated perivascular fibroblasts, and interstitial fibroblasts, are located medial to the somite. Interestingly, progenitors with initial dorsal projections predominantly generated type I cells (13/14, 93%), whereas cells projecting ventrally were more likely to give rise to type II cells (11/19, 58%) than type I cells (3/19, 16%) and sometimes generated cells of both types (5/19, 26%) (Fig 6E-F). It is important to note that ventral projecting progenitors at 24 hpf never generated dorsal fin mesenchymal cells and all type I cells generated by this group were DLAV fibroblasts found along the ventral side of the DLAV (Fig 6F). Together, our results suggest that the dorsal sclerotome domain contains a mixture of unipotent and bipotent progenitors at 24 hpf, and their migration direction likely restricts the types of fibroblasts they differentiate into.

**Fig 6.**
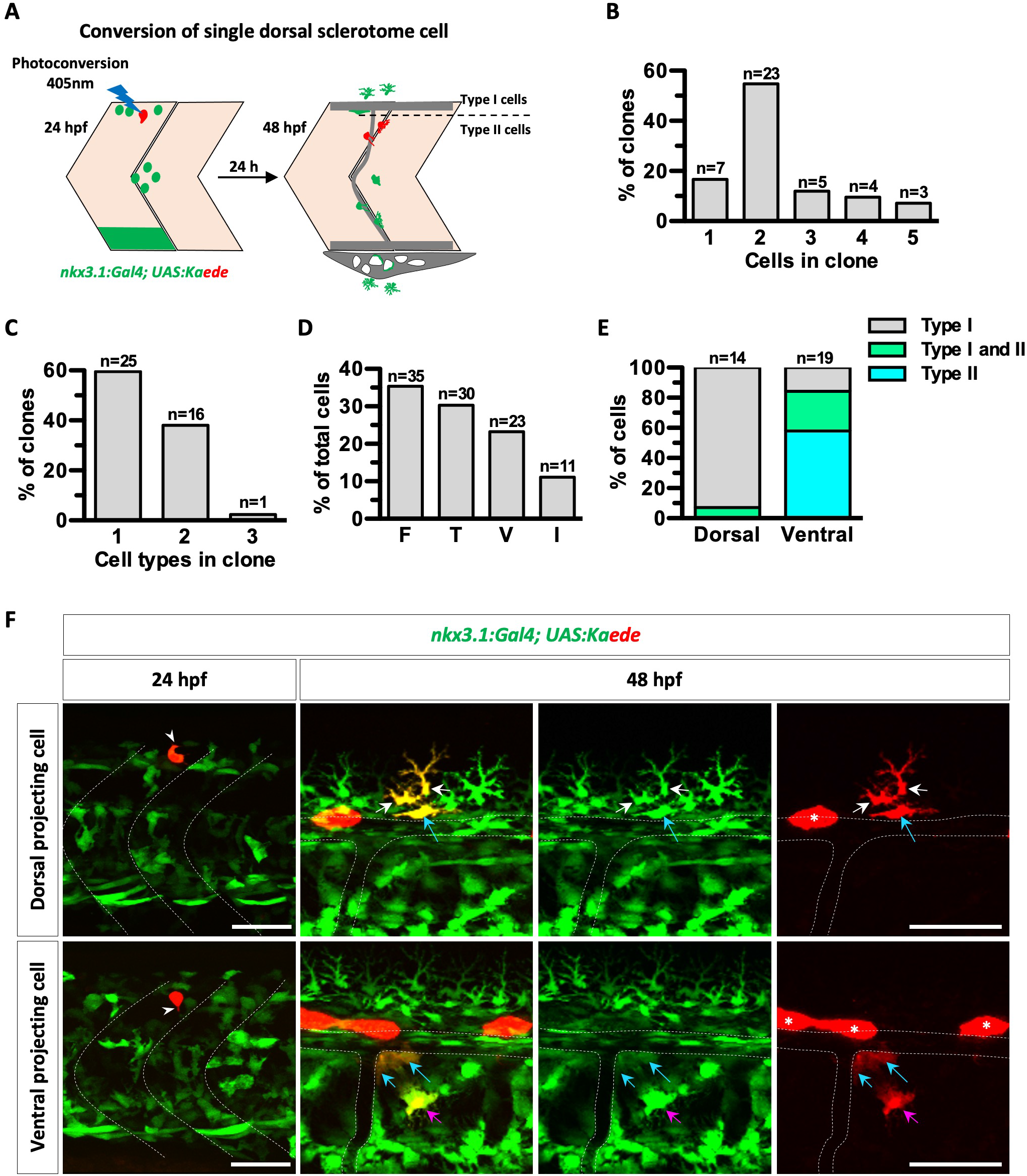
Clonal analysis of the dorsal sclerotome. (A) Schematic of clonal analysis. A single dorsal sclerotome progenitor in *nkx3.1*^*Kaede*^ embryos was photoconverted at 24 hpf and imaged at 48 hpf to determine clone composition. Dorsal sclerotome-derived cells were subdivided into two groups based on their dorsal-ventral positions: the more dorsal type I cells include dorsal fin mesenchymal cells and DLAV fibroblasts (above the dotted line), while the more ventral type II cells include tenocytes, perivascular fibroblasts, and interstitial fibroblasts (below the dotted line). (B) Quantification of the clone size. (C) Quantification of fibroblast subtypes in each clone. (D) Distribution of different fibroblast subtypes from all decedents of traced cells. F, fin mesenchymal cells; T, tenocytes; V, blood vessel associated fibroblasts; I, interstitial fibroblasts. (E) Quantification of the clone composition. Only photoconverted cells showing polarized cell projections at 24 hpf were graphed. *n*=14 (with dorsal projection) and 19 (with ventral projection) cells. (F) Examples of clonal analysis of dorsal sclerotome progenitors in *nkx3.1*^*Kaede*^ embryos at 24 and 48 hpf. In the top panel (type I cell lineage), the photoconverted cell showing dorsal projections (arrowhead) generated two dorsal fin mesenchymal cells (white arrows) and one DLAV fibroblast (long cyan arrows). In the bottom panel (type II cell lineage), the photoconverted cell with ventral cell projections (arrowhead) generated one DLAV fibroblast (long cyan arrows), one perivascular fibroblast (cyan arrows), and one tenocyte (magenta arrows). The outline of the vasculature is denoted by dotted lines. *n*=42 photoconverted cells from 40 embryos. Scale bars: 50 μm.

### Fate of ventral sclerotome progenitors is biased by the position along the migratory path

In posterior somites, some ventral sclerotome progenitors migrate ventrally towards the CVP region to generate SRCs and ventral fin mesenchymal cells. To visualize this dynamic process, we performed time-lapse imaging of *kdrl:EGFP; nkx3.1*^*NTR-mCherry*^ embryos from 25 to 50 hpf (Video S1 and Fig S3). At 25 hpf, some migrating mCherry^+^ progenitors were found ventral to the somites. These cells migrated ventrally towards the fin fold as more mCherry^+^ cells emerged from the ventral somite and joined the migration. To better describe the position of migrating cells, we referred to cells at the front edge of the migration as “leading cells”, while all cells behind were called “lagging cells”. We observed a stereotypical migration pattern whereby mCherry^+^ cells “marched” ventrally in an orderly fashion, and individual cells did not pass other cells that were ahead. This orderly migration was largely maintained, even during cell division. As a result, descendants of leading cells remained at the front of the migration throughout the movie and formed ventral fin mesenchymal cells, whereas descendants of lagging cells mostly generated SRCs associated with the CVP (Video S1 and Fig S3). Interestingly, the cardinal vein underwent similar ventral expansion to form the CVP during the same time window. Lagging cells appeared to constantly interact with the forming CVP, while leading cells remained ahead of the vascular sprouts (Video S1 and Fig S3). Together, our results suggest that the fate of ventral sclerotome progenitors is biased by their position along the migratory path, potentially due to their interactions with the local environment.

To further compare the lineage potential between the leading and lagging cells, we performed single-cell clonal analysis in *nkx3.1*^*Kaede*^ fish (Fig 7A). Single ventrally migrating Kaede^green^ progenitors between somites 21-25 were photoconverted to Kaede^red^ at 24 hpf, and their descendants were imaged after 24 hours. Similar to our time-lapse movies, the majority of photoconverted leading cells generated only ventral fin mesenchymal cells (17/20, 85%), but a small number gave rise to both SRCs and ventral fin mesenchymal cells (3/20, 15%) (Fig 7B-C). Interestingly, leading cells never generated SRCs only. By contrast, 41% of lagging cells generated only SRCs (9/22), 36% generated both ventral fin mesenchymal cells and SRCs (8/22), and 23% produced only ventral fin mesenchymal cells (5/22) (Fig 7B-C). Combining time-lapse imaging and single-cell tracing, our results suggest that the relative position along the migratory path biases the fate of these sclerotome-derived cells.

**Fig 7.**
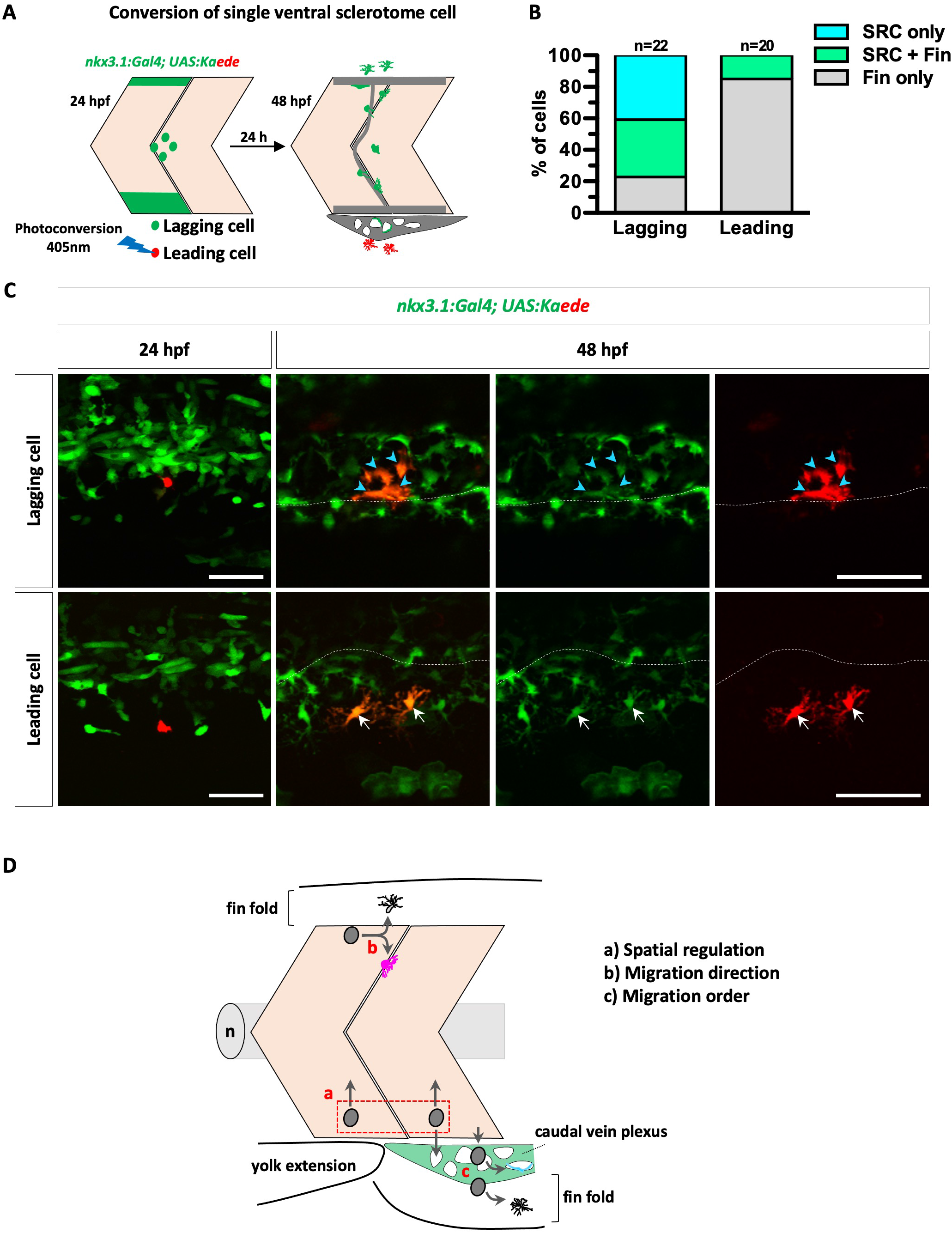
Clonal analysis of the ventral sclerotome. (A) Schematic of clonal analysis. A single ventral sclerotome progenitor in *nkx3.1*^*Kaede*^ embryos was photoconverted at 24 hpf and imaged after 24 hours to determine clone composition. Cells were classified as leading or lagging cells based on their initial position at 24 hpf, where cells at the front of the migration path were defined as leading cells. (B) Quantification of the clone composition. (C) Examples of cell tracing at 24 and 48 hpf. A lagging cell (top panel) generated four SRCs (arrowheads) dorsal to the ventral edge of the CVP (dotted lines). A leading cell (bottom panel) generated two fin mesenchymal cells (arrows). *n*=42 photoconverted cells from 38 embryos. (D) Three mechanisms contribute to the diversification of fibroblast subtypes from sclerotome progenitors in zebrafish. The fate of sclerotome progenitors is biased by their dorsal-ventral or anterior-posterior positions (a, spatial regulation), migration direction (b), and migration order (c). n, notochord. Scale bars: 50 μm.

## DISCUSSION

Our work establishes the sclerotome as the embryonic source of different connective tissue cells in zebrafish. First, the sclerotome gives rise to distinct fibroblasts in the trunk. Second, the sclerotome contributes to proper development of the axial skeleton. Third, the sclerotome contains a mixture of unipotent and bipotent progenitors and their differentiation is biased by their axial positions and migratory paths.

### The sclerotome is the embryonic source of trunk fibroblasts

The developmental potential of the sclerotome in zebrafish has been first explored with vital dye injections (Morin-Kensicki and Eisen, 1997). Using a classic sclerotome marker *nkx3.1* (Schneider et al., 2000), we have previously developed the *nkx3.1:Gal4* line in zebrafish to genetically label sclerotome progenitors and their descendants (Ma et al., 2018). Here, we provide additional evidence that *nkx3.1:Gal4* labels cells of the sclerotome lineage. First, the *nkx3.1*^*NTR-mCherry*^ reporter co-expresses known sclerotome markers such as *versican b* (Landolt et al., 1995), *foxc1b* (Topczewska et al., 2001), and *twist1b* (Germanguz et al., 2007; Yeo et al., 2009). Second, lineage tracing of *nkx3.1*-derived cells shows extensive labeling of cells associated with the axial skeleton, consistent with previous cell tracing experiments (Morin-Kensicki et al., 2002). Interestingly, the *nkx3.1*^*NTR-mCherry*^ reporter also labels some muscles adjacent to the dorsal and ventral sclerotome domains. This observation is in agreement with previous single-cell lineage analysis of the presumptive sclerotome in the ventral somite where a sub-population of these cells gives rise to both sclerotome derivatives and muscles (Morin-Kensicki and Eisen, 1997). We speculate that some bipotent somitic cells might transiently express *nkx3.1*, before committing to either the myotome or the sclerotome lineage. This idea is supported by the observation that progenitors at developmental branch points sometimes express markers characteristic of multiple fates (Farrell et al., 2018).

Our previous studies (Ma et al., 2018; Rajan et al., 2020) and current work provide strong evidence that the sclerotome is the embryonic source of multiple types of fibroblasts in the trunk of zebrafish. Based on marker expression, anatomical location, cell morphology, and function, we define at least four fibroblast subtypes derived from the sclerotome: tenocytes, fin mesenchymal cells, blood vessel associated fibroblasts, and interstitial fibroblasts. Tenocytes express classic tendon markers *scxa* and *tnmd* (Chen and Galloway, 2014; Ma et al., 2018). Their cell bodies are positioned medially along the myotendinous junction, while extending long cellular processes into the intersomitic space (Ma et al., 2018; Subramanian et al., 2018). Tenocytes stabilize muscle attachment likely by secreting a number of ECM proteins (Ma et al., 2018; Subramanian and Schilling, 2014). Fin mesenchymal cells, on the other hand, are marked by the expression of ECM genes *hmcn2* and *fbln1*. They are located adjacent to apical epidermal cells in the fin fold with distally extended tree-like processes. Recent work has shown that fin mesenchymal cells guide fin formation by aligning collagen fibers of actinotrichia (Kuroda et al., 2020). Genetic ablation of fin mesenchymal cells in the pectoral and median fin folds results in collapse of the respective fins (Lalonde and Akimenko, 2018). Although blood vessel associated fibroblasts are labeled by pan-fibroblast markers similar as other fibroblast subtypes in the trunk, they are characterized by their close association with the vasculature. Interestingly, their morphology differs depending on the vascular bed. For example, ISV-associated perivascular fibroblasts are globular, whereas DLAV fibroblasts are more elongated. We have previously shown that perivascular fibroblasts play dual roles in stabilizing nascent ISVs and functioning as pericyte precursors (Rajan et al., 2020). It is likely that other blood vessel associated fibroblasts play similar roles in stabilizing the corresponding blood vessels. Interestingly, stromal reticular cells closely interact with CVP, which functions as an early hematopoietic niche during development (Wattrus and Zon, 2018). Indeed, recent work has shown that defects in SRC maturation result in a compromised CVP niche that fails to support hematopoietic stem cell (HSC) maintenance and expansion (Murayama et al., 2015). Therefore, SRCs might not only provide structural support to the CVP, but also modulate the microenvironment required for HSC differentiation. Despite extensive labeling of blood vessel associated fibroblasts, our lineage analysis shows that sclerotome progenitors do not contribute to the vasculature. This result suggests that the sclerotome domain defined by *nkx3.1* is distinct from the endotome, a sub-compartment of the somite that has been shown to contain endothelial precursors (Nguyen et al., 2014; Sahai-Hernandez et al., 2020). Compared to other fibroblast subtypes, interstitial fibroblasts are the least well-characterized. Based on their distribution around the notochord and spinal cord, it is conceivable that interstitial fibroblasts give rise to osteoblasts required for vertebral column development. This is consistent with our observation that long-term cell tracing of the *nkx3.1* lineage results in extensive labeling of cells associated with the vertebral bodies, neural and hemal arches. Together, our results suggest that the sclerotome generates distinct fibroblast subtypes to support a variety of tissues, including muscles, fin folds and blood vessels.

### The sclerotome contributes to axial skeleton development in zebrafish

In amniotes, the sclerotome is the embryonic source of the vertebrae, structural units of the vertebral column (Christ et al., 2004). In teleosts, each vertebra is composed of three elements: vertebral body (centrum), neural, and hemal arches. In contrast to amniotes, studies in zebrafish and salmon have suggested that the centrum is formed by notochord sheath cells instead of the sclerotome (Fleming et al., 2004; Grotmol et al., 2003; Lleras Forero et al., 2018; Pogoda et al., 2018; Wang et al., 2013; Wopat et al., 2018). However, how the sclerotome contributes to the development of the axial skeleton in teleosts remains unclear. Our long-term lineage tracing experiments demonstrate that cells derived from the sclerotome lineage are closely associated with all three skeletal elements of the vertebrae. Partial depletion of sclerotome-derived cells results in variable defects in neural and hemal arches, while centra are largely normal. This result is consistent with the current model, in which the centrum forms independently of the somite (Lleras Forero et al., 2018; Wopat et al., 2018). However, it is not clear why sclerotome ablation leads to a stronger defect in neural arches than hemal arches, despite similar labeling of sclerotome-derived cells on both structures. There are three non-mutually exclusive explanations. First, severe loss of hemal arches might be detrimental to survival; therefore, only fish with mild phenotypes can survive to 3 wpf (time of analysis). Second, the precursors of hemal arches might have a greater capacity to regenerate after ablation than those of neural arches. Finally, there might be an alternative non-sclerotome source that contributes to the formation of hemal arches redundantly. Interestingly, our results in zebrafish show remarkable parallel to previous observations in medaka. In medaka, sclerotome-derived cells marked by *twist:GFP* are similarly enriched in the intervertebral regions, and are thought to be the source of osteoblasts to generate different skeletal elements (Inohaya et al., 2007). Morpholino knockdown of key sclerotome genes, including *pax1, pax9*, and *twist*, results in the loss of neural arches with largely normal centra (Mise et al., 2008; Yasutake et al., 2004). Together, these results suggest that the sclerotome contributes to proper development of the axial skeleton in teleosts.

### The developmental origin of fin mesenchymal cells

There has been considerable debate regarding where fin mesenchymal cells originate during fish and amphibian development. Lineage tracing studies in zebrafish suggest that the paraxial mesoderm, specifically the dermomyotome, but not the sclerotome, is the embryonic source of fin mesenchymal cells (Lee et al., 2013). Their conclusions contrast with our findings in two aspects. First, using the *pax3a:EGFP* line as a marker for the dermomyotome, Lee et al. (2013) show extensive contribution of EGFP^+^ fin mesenchymal cells throughout the entire major lobe of the fin fold, including the caudal fin. By contrast, we find that a different dermomyotome reporter, *pax7b*^*EGFP*^ (Pipalia et al., 2016), contributes minimally to fin mesenchymal cells in the fin fold. This discrepancy is likely due to the different expression dynamics of *pax3a* and *pax7b*, despite both being dermomyotome markers. It has been shown that *pax3a* is initially broadly expressed in the entire somite before becoming restricted to the anterior somites (precursors of the dermomyotome) (Feng et al., 2006; Hammond et al., 2007). However, *pax7b* is expressed several hours after *pax3a* and is restricted to the anterior somites (Feng et al., 2006; Hammond et al., 2007; Minchin and Hughes, 2008). Therefore, the *pax3a:EGFP* line may not be as specific as *pax7b*^*EGFP*^ in labeling the dermomyotome. Second, Lee et al. (2013) use the *ola-twist1:Gal4* transgenic line, in which Gal4 expression is driven by the medaka (*Oryzias latipes*) *twist1* promoter, to rule out sclerotome contribution. However, our results using the *nkx3.1*^*NTR-mCherry*^ reporter indicate that the sclerotome contributes to most fin mesenchymal cells in both the dorsal and ventral regions of the major lobe. Interestingly, we have previously shown that the expression of a similar *ola-twist1:EGFP* reporter is restricted to sclerotome-derived cells around the notochord, but is absent in both dorsal and ventral sclerotome domains (Ma et al., 2018). This observation explains why the *ola-twist1:Gal4* line fails to label any fin mesenchymal cells (Lee et al., 2013). Together, our work suggests an updated model where fin mesenchymal cells are generated by multiple lineages, depending on their locations. Fin mesenchymal cells in the dorsal and ventral fin fold have a sclerotome origin, those in the caudal fin originate from a non-dermomyotome and non-sclerotome compartment of the somite, while fin mesenchymal cells in the minor lobe of the fin fold have an unidentified embryonic source.

### Mechanisms of fibroblast diversification

A fundamental question in developmental biology is how progenitors in the embryo diversify into distinct differentiated cell types. Using *nkx3.1*-based transgenic reporters, our work reveals three key features of cell fate diversification from the sclerotome lineage (Fig 7D). First, sclerotome progenitors display regional differences in their differentiation potential along the dorsal-ventral and anterior-posterior positions. Dorsally located fibroblasts, such as dorsal fin mesenchymal cells and DLAV fibroblasts, are derived from the dorsal sclerotome, whereas ventrally located fibroblasts, such as stromal reticular cells and ventral fin mesenchymal cells, are generated by the ventral sclerotome. Similarly, the ventral sclerotome in anterior somites (before the end of the yolk extension) generates only fibroblasts in the trunk, whereas the ventral sclerotome in posterior somites contributes to stromal reticular cells and fin mesenchymal cells in addition to trunk fibroblasts. Second, fate of sclerotome progenitors is biased by their migratory path. For dorsal sclerotome progenitors, dorsal migration limits the cell to become either dorsal fin mesenchymal cells or DLAV fibroblasts, while ventral migration leads to the differentiation of predominately tenocytes or perivascular fibroblasts. Third, the relative order of migrating sclerotome progenitors highly correlates with their differentiation potential. Leading cells are more likely to become ventral fin mesenchymal cells, while lagging cells tend to differentiate into stromal reticular cells. The combination of these three mechanisms (spatial regulation, migration direction, and migration order) (Fig 7D) likely contributes to the generation of different fibroblast subtypes in a stereotypical manner (Ma et al., 2018; Rajan et al., 2020).

How does a sclerotome progenitor decide which cell type to differentiate into? Photoconversion-based clonal analysis reveals that single sclerotome progenitors predominantly generate 1-2 different fibroblast subtypes. This result suggests that the sclerotome contains a mixture of unipotent and bipotent progenitors prior to migration. The presence of bipotent progenitors suggests that individual sclerotome cells are plastic and are not fate-restricted to become a specific fibroblast subtype. We speculate that the local microenvironment plays a key role in regulating the migration and subsequent differentiation of sclerotome-derived cells. In the trunk region, the migration of sclerotome-derived cells depends on Sonic Hedgehog (Shh) from the notochord and floor plate (Ma et al., 2018). Inhibition of Shh signaling results in the loss of tenocytes between muscle boundaries (Ma et al., 2018). Similarly, Slit-Robo signaling has been shown to mediate the polarization and migration of fin mesenchymal cells in the peripheral fin fold (Mahabaleshwar et al., 2022). Cell type diversification of the sclerotome is remarkably similar to that of the neural crest lineage. Previous studies have shown that migrating neural crest cells are multipotent (Baggiolini et al., 2015), and the ultimate cell fate is determined by their axial positions as well as the migration timing and direction (Rocha et al., 2020).

In summary, we demonstrate that the sclerotome is the embryonic source of multiple fibroblast subtypes and contributes to axial skeletal development in zebrafish. Our work shows that the sclerotome is an excellent model for studying cell type diversification during embryonic development. The combination of positional and migratory information as well as the local microenvironment likely contributes to the differentiation trajectory of sclerotome progenitors. Future work using scRNA-seq may further reveal the heterogeneity of sclerotome-derived cells and identify key gene regulatory networks underlying the lineage progression of sclerotome progenitors.

## MATERIALS AND METHODS

### Ethics statement

All animal experiments were conducted in accordance with the principles outlined in the current Guidelines of the Canadian Council on Animal Care. All protocols were approved by the Animal Care Committee of the University of Calgary (#AC17-0128 and #AC21-0102).

### Zebrafish strains

Zebrafish strains used in this study were maintained and raised under standard conditions. The following transgenic strains were used in this study: *Gt(:gSAIz-GAL4FFD)nkgsaizGFFD164A* (*pax7b:Gal4FF*) (Pipalia et al., 2016), *TgBAC(col1a2:Gal4)ca102* (Ma et al., 2018; Sharma et al., 2019), *TgBAC(col1a2:GFP)ca103* (Ma et al., 2018), *Tg(kdrl:EGFP)la163* (Choi et al., 2007), *Tg(kdrl:mCherry)ci5* (Proulx et al., 2010), *TgBAC(nkx3.1:Gal4)ca101* (Ma et al., 2018), *Tg(UAS:EGFP)nkuasgfp1a* (Asakawa et al., 2008), *Tg(UAS:Cre-ERT2)ca105* (Sharma et al., 2019), *Tg(−3.5ubb:LOXP-EGFP-LOXP-mCherry)cz1701* (*ubi:Switch*) (Mosimann et al., 2011), *Tg(UAS:Kaede)s1999t* (Davison et al., 2007), and *Tg(UAS:NTR-mCherry)c264* (Davison et al., 2007). The mosaic *col1a2:Gal4; UAS:Kaede* line was maintained by selecting embryos with more mosaic Kaede expression.

### In situ hybridization and immunohistochemistry

Whole-mount in situ hybridization and immunohistochemistry were performed according to previously established protocols. We used the following RNA probes in this study: *cilp, col5a1, fbln1, foxc1b, fras1, gal4, hmcn2, kaede, ntr-mCherry, scxa, tgfbi, tnmd, twist1b, vcanb* (*versican b*). Double fluorescent in situ hybridizations were performed using different combinations of digoxigenin (DIG) or dinitrophenyl (DNP) labeled probes. For immunohistochemistry, rabbit polyclonal antibody to RFP (1:1000, MBL, PM005) and chicken antibody to GFP (1:250, Aves Labs, GFP-1020) were used. For fluorescent detection of antibody labeling, the appropriate Alexa Fluor-568 or Alexa Fluor-647 conjugated secondary antibodies (1:500, Thermo Fisher) were used.

### Kaede photoconversion

Photoconversion was carried out using a 405 nm laser and a 20x objective on an Olympus FV1200 confocal microscope. At the appropriate stage, Kaede-expressing embryos were anesthetized using tricaine and mounted in 0.8% low-melting-point agarose. The duration and intensity of laser required for complete green-to-red conversion depended on the Kaede expression level and the size of the target region. For cell tracing of the sclerotome domains, a circular region of 20×20 pixels was converted using 1.5% 405 nm laser for 4 seconds. For single-cell clonal analysis of sclerotome progenitors, typically a circular region of 10×10 pixels was converted using 1.5% 405 nm laser for 4 seconds. After photoconversion, embryos were confirmed for the expression of Kaede^red^ and recovered in E3 fish water until the necessary stages.

### Cre-mediated lineage tracing

*nkx3.1:Gal4; UAS:Cre-ERT2; ubi:Switch* embryos were treated with 10 µM 4-hydroxytamoxifen (4OHT) from 1 to 2 dpf. After treatment, embryos were washed three times and recovered in fish water for analysis at the appropriate stages.

### Cell ablation

NTR-mCherry^+^ embryos and NTR-mCherry^-^ sibling controls at the appropriate stage were treated with metronidazole (MTZ, Sigma Millipore) at a concentration of 5 mM in fish water. Embryos were incubated in MTZ solution for at least 24 hours to achieve complete ablation. Fish were then washed and recovered in fish water until the desired stage for analysis.

### Bone and cartilage staining

Bone and cartilage staining was performed following a modified version of a previously described protocol (Walker and Kimmel, 2007). Briefly, larvae at 3 wpf were euthanized in 0.4% tricaine and fixed in 4% formaldehyde in PBS overnight at 4 °C. Fixed larvae were washed in a solution containing 100 mM Tris pH 7.5 and 10 mM MgCl_2_, then stained overnight in 0.04% alcian blue solution. The larvae were then washed with decreasing concentrations of ethanol diluted in 100 mM Tris pH 7.5 (80%, 50%, 25%) and bleached in a 3% H_2_O_2_/0.5% KOH solution until all pigments were lost. They were washed twice with 25% glycerol/0.1% KOH and stained in the same solution with 0.01% alizarin red for 1 hour. Finally, larvae were washed twice in 50% glycerol/0.1% KOH and switched to 100% glycerol for imaging.

### Calcein staining

Bones can be fluorescently labelled in live larvae using the vital dye, calcein. Larvae at 17 dpf were placed in a 100 mm petri dish in a 0.2% calcein solution for 15 – 30 minutes at room temperature. They were then washed at least six times with water from zebrafish vivarium. The larvae were left for 2 – 3 hours prior to imaging to allow excess calcein to pass through their digestive tract. Fish were anesthetized using tricaine and mounted in 0.8% low-melting-point agarose for live imaging. To obtain transverse views, fish were fixed in 4% formaldehyde and sectioned for imaging.

### Vibratome sectioning

Juvenile and adult zebrafish were fixed in 4% paraformaldehyde (PFA) with 1% dimethyl sulfoxide (DMSO) overnight at 4 °C. Fixed fish were washed in PBS and decalcified in 0.5 M EDTA at room temperature for 3 days. Following decalcification, the dissected trunks were embedded in 20% gelatin and sectioned using a Leica VT1000S vibratome to obtain slices at 150 μm thick. The sections were mounted on slides in 50% glycerol for confocal imaging. Confocal tile scans were stitched together using Olympus Fluoview software to generate a full view of the transverse section.

### Time-lapse imaging and cell tracking

Time-lapse imaging of zebrafish embryos was carried out using an Olympus FV1200 confocal microscope. At the appropriate stage, fish were anesthetized using tricaine and mounted in 0.8% low-melting-point agarose. To keep the agarose hydrated and minimize the development of pigments, as well as minimize movement of the fish, a small volume of E3 water with phenylthiourea and tricaine was carefully flooded around the agarose. Z-stack images of the region of interest were collected at regular intervals (6-9 mins) for up to 25 hours. Images were then compiled into movies using Olympus Fluoview software. Cells from each movie were manually tracked using the Fiji software (Schindelin et al., 2012).

### Statistical analysis

All graphs were generated using the GraphPad Prism software. Data were plotted as mean±SEM. The significance between two samples was calculated using Mann-Whitney *U* test. p values: p>0.05 (not significant); p<0.05 (*); p<0.01 (**); p<0.001 (***); p<0.0001 (****).

## ACKNOWLEDGMENTS

We thank the zebrafish community for providing probes and reagents; Simon Hughes for sharing the *pax7b*^*EGFP*^ line; Sarah Childs for sharing reagents, transgenic lines and providing critical input on this project; members of the Childs and Huang laboratories for discussions; Arsheen Rajan for critical comments on the manuscript.

## SUPPLEMENTAL FIGURES

**Fig S1.**
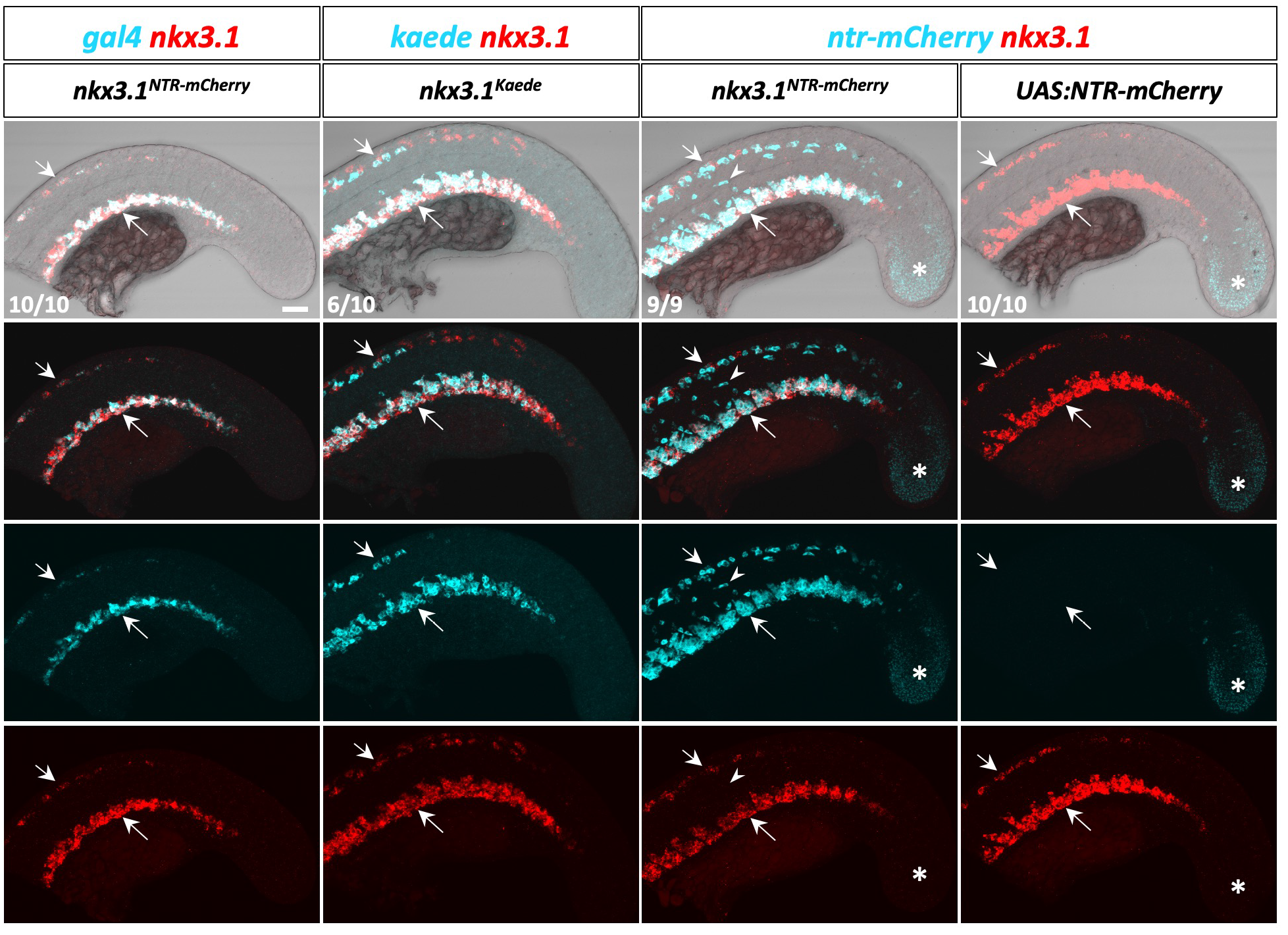
Characterization of the sclerotome reporter. Double fluorescent in situ staining of *nkx3.1* (red) together with *gal4, kaede*, or *ntr-mcherry* (cyan) was performed in *nkx3.1*^*NTR-mCherry*^, *nkx3.1*^*Kaede*^, or *UAS:NTR-mCherry* embryos at 20 hpf. *gal4* expression in *nkx3.1*^*NTR-mCherry*^ embryos accurately recapitulated endogenous *nkx3.1* expression in both dorsal and ventral sclerotome domains (short and long arrows, respectively). *kaede* expression in *nkx3.1*^*Kaede*^ embryos was largely consistent with *nkx3.1* expression, whereas *nkx3.1*^*NTR-mCherry*^ embryos showed ectopic *ntr-mcherry* expression in somites (arrowheads). Note that *UAS:NTR-mCherry* but not *UAS:Kaede* embryos also showed low Gal4-independent expression in the tail bud (asterisks). Representative images are shown with *n* numbers for each staining. Both the merged and individual channels are shown. Scale bar: 50 μm.

**Fig S2.**
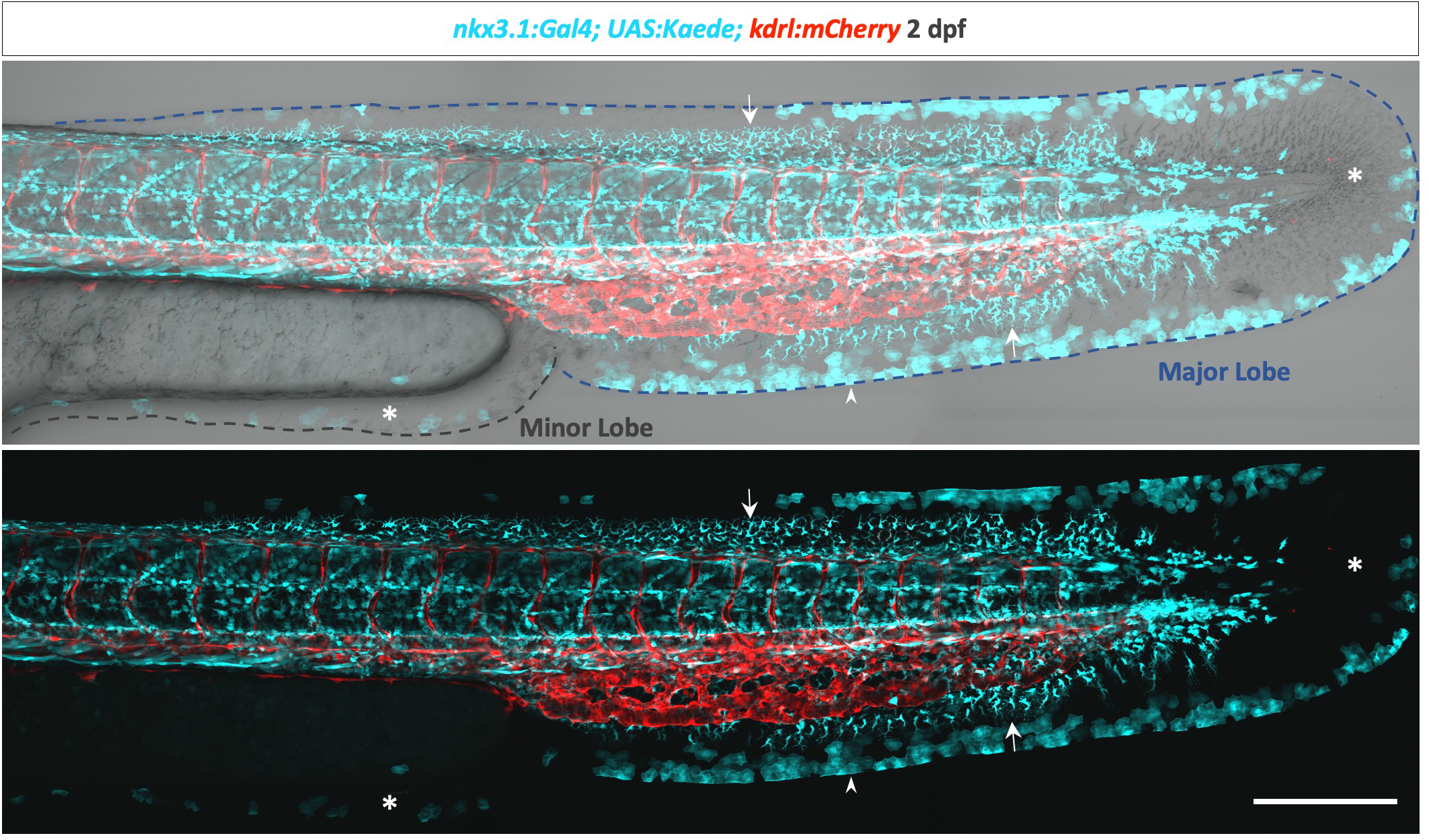
The sclerotome generates fin mesenchymal cells in the dorsal and ventral fin folds. A tiled confocal image of *nkx3.1*^*Kaede*^; *kdrl:mCherry* embryo at 2 dpf showing Kaede^+^ sclerotome-derived cells (cyan) throughout the trunk of the embryo. Kaede^+^ fin mesenchymal cells were present in both dorsal and ventral fin folds (arrows), but absent from either the caudal fin or the minor lobe (asterisks). Arrowheads indicate mCherry expression in skin cells at the edge of the fin fold. The major and minor lobes of the fin fold are outlined by dashed lines. *n*=5. Scale bar: 200 μm.

**Fig S3.**
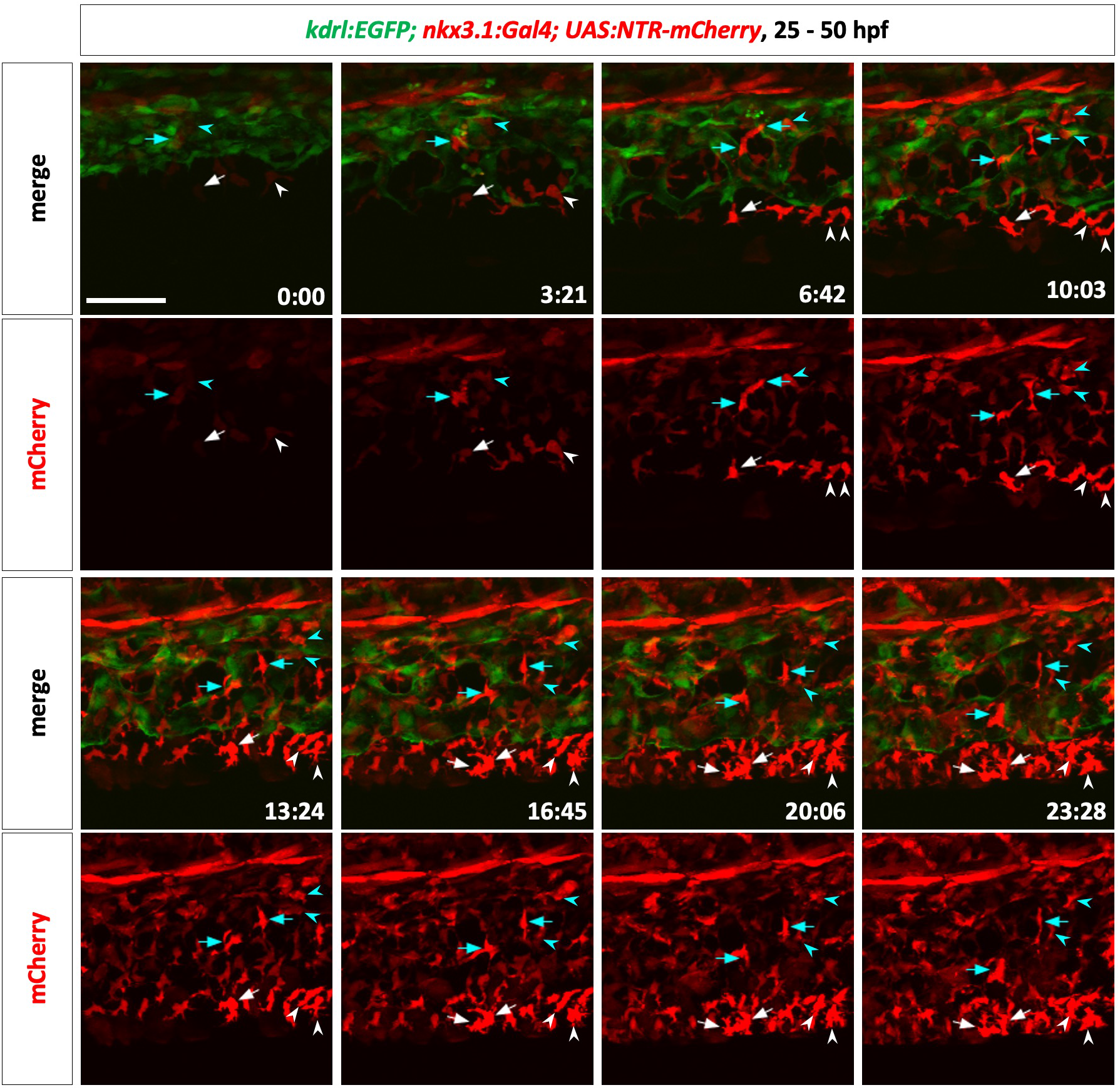
Time-lapse imaging of migration of ventral sclerotome progenitors. Snapshots of a *kdrl:EGFP; nkx3.1*^*NTR-mCherry*^ embryo between 25 and 50 hpf are shown with timestamps (hh:mm). The migration and division of two representative lagging cells (cyan arrows and arrowheads) and two leading cells (white arrows and arrowheads) were traced throughout the movie. Daughter cells are indicated by arrows or arrowheads in the same color. The lagging cell never migrated ahead of the leading cell and generated 2 SRCs. The leading cell remained at the migration front ahead of the lagging cell and generated 2 ventral fin mesenchymal cells. *n*=7. Scale bar: 50 μm.

**Video S1. Time-lapse imaging of migration of ventral sclerotome progenitors**. *kdrl:EGFP; nkx3.1*^*NTR-mCherry*^ embryos were imaged from 25 to 50 hpf with timestamps (hh:mm). The migration and division of representative lagging (cyan arrows) and leading (white arrows) cells were traced throughout the movie. Daughter cells are indicated by arrows in the same color. The lagging cell never migrated ahead of the leading cell and generated 2 SRCs. The leading cell remained at the migration front ahead of the lagging cell and generated 2 ventral fin mesenchymal cells. Snapshots of this video are shown in Fig S3. *n*=7. Scale bar: 50 µm.

